# Plaque-associated endogenous IgG and its impact on immunohistochemical detection of mouse monoclonal IgG antibodies in mouse models of Alzheimer’s disease

**DOI:** 10.1101/2024.04.25.591057

**Authors:** Shogo Ito, Kenta Yamauchi, Hiroshi Hama, Masato Koike, Atsushi Miyawaki, Hiroyuki Hioki

## Abstract

Experimental studies for Alzheimer’s disease (AD) have largely depended on transgenic mice with β-amyloidosis. Here, we report plaque-associated endogenous immunoglobulin G (PA-IgG) and its impact on indirect immunohistochemical detection of mouse monoclonal IgG antibodies (Ms monoclonal IgG Abs) in the brain of AD mouse models. Immunostaining for Ms IgG in AD mouse models demonstrated endogenous IgG in the brain parenchyma accumulated on microglia associated with amyloid β (Aβ) plaques and/or Aβ plaques themselves. This PA-IgG caused robust off-target binding of secondary Abs against Ms IgG (H+L) in indirect immunohistochemistry using Ms monoclonal IgG Abs. Blocking with Fab fragments of anti-Ms IgG (H+L) Ab was not effective against off-target binding. Unexpectedly, we found that secondary Abs that specifically recognize Ms IgG1, 2a, 2b, and 3 did not cause off-target binding on frozen brain sections of *App^NL-G-F/NL-G-F^*mice, and enabled specific labeling of Ms monoclonal IgG Abs in the AD mouse model brains. We further demonstrated that indirect detection with a conventional secondary Ab against Ms IgG (H+L) Ab could lead to erroneous conclusions regarding Aβ plaque burden and phosphorylated tau accumulation in *App^NL-G-F/NL-G-F^*mice, and the use of Ms IgG subclass specific secondary Abs allowed to avoid the inevitable impediment caused by the endogenous IgG accumulation. Specific indirect detection of Ms monoclonal IgG Abs in AD mouse models by the use of secondary Abs against Ms IgG subclass would accelerate AD research by expanding the choice of Abs available for histochemical analysis in AD studies.

## Introduction

Alzheimer’s disease (AD) is the most prevalent neurodegenerative disease that accounts for 60 – 80% of all dementia cases^1^. AD patients present with progressive cognitive decline, and cortical and hippocampal atrophy^2^. Neuritic plaques and neurofibrillary tangles (NFTs) represents the major histopathological hallmarks of AD^2,3^. Neuritic plaques comprise extracellular deposition of amyloid β (Aβ) peptides, while NFTs are intracellular accumulation of phosphorylated and misfolded tau proteins^4^. Reactive gliosis, abnormal proliferation and morphology of astrocytes and microglia, is another histopathological feature of AD^5–7^. Currently, no causal therapy is developed for AD that halts or reverses the progression of cognitive impairments. The amyloid hypothesis that Aβ plaque accumulation in the brain is the primary cause of AD^8–10^ is supported by genetic and pathological evidence^10^.

Experimental studies for AD have largely depended on transgenic mice with β-amyloidosis in their brains^11^. Given mutations in *amyloid precursor protein* (*App*) and its cleaving enzyme *presenilin* (*Psen*) *1* and *2* in early-onset familial AD (FAD), numerous transgenic mice that express and/or overexpress APP or APP/PSEN1/2 with the mutations have been generated for mouse models of AD^12,13^. These mice overproduce pathogenic Aβ peptides and exhibit several key features of AD, such as age-dependent impairments in learning and memory, synaptic deficits, aberrant neuronal network activity, formation of Aβ plaques, and neuroinflammatory responses of astrocytes and microglia^12,14,15^. Recently, AD mouse models utilizing knock-in strategies have been generated to introduce *FAD* mutations into the endogenous mouse APP gene (*App*-knockin (KI) mouse models), avoiding overexpression of APP gene that could lead to artificial effects of N- and C-terminal fragments of APP protein^14,16^.

Mouse monoclonal immunoglobulin G (Ms monoclonal IgG) antibodies (Abs) have been used in histological studies and pathological diagnosis by combing with the indirect immunohistochemical method. Indirect immunohistochemistry (IHC) employs secondary Abs conjugated to enzymes, haptens, or fluorophores that recognize unlabeled primary Abs. The indirect method is often preferred to the direct method that uses directly conjugated primary Abs. Indirect detection demonstrates higher sensitivity than the direct method^17,18^. Moreover, a single secondary Ab in indirect IHC can be used to detect a wide range of primary Abs generated in a given species. Monoclonal Abs are more specific and reproducible compared to polyclonal Abs which are composed of a heterogenous mixture of Abs that recognize the same antigen. Monoclonal Abs were first generated in mice by fusing their antibody-producing splenocytes with myeloma cells^19^. Over the past decades, thousands of monoclonal Abs have been generated from hybridomas producing mouse IgG (Ms IgG), and utilized for various experimental and clinical studies. There are five IgG subclasses, IgG1, 2a, 2b, 2c, and 3, in the mouse^20–22^. The variety of Ms IgG has been used for multiple labeling of Ms monoclonal IgG Abs, generating secondary Abs that specifically recognize each Ms IgG subclass. Indirect IHC detection of Ms monoclonal IgG Abs in AD mouse models is indispensable for experimental research for AD such as the evaluation of developing therapies for the diseases^23^. Ms monoclonal IgG Abs against Aβ and phosphorylated tau, such 4G8, 6E10, and AT8, have been golden standards for experimental studies and pathological diagnosis for AD^24–26^. However, endogenous IgG accumulation in AD mouse models^27–29^ could confound specific detection of Ms monoclonal IgG Abs by causing off-target binding of secondary Abs that recognize IgG molecules of mice.

Here, we show endogenous IgG accumulation associated with Aβ plaques, namely plaque-associated endogenous IgG (PA-IgG), and its impact on indirect IHC with Ms monoclonal IgG Abs in AD mouse models. The application of fluorophore-conjugated Abs against Ms IgG heavy and light chains [IgG (H+L)] to frozen brain sections of *App^NL-F/NL-F^* and *App^NL-G-F/NL-G-F^*mice demonstrated robust accumulation of endogenous IgG on microglial cells associated with Aβ plaques (PAM) and/or Aβ plaques themselves. PA-IgG engaged anti-Ms IgG (H+L) secondary Abs on brain tissues of *App^NL-G-F^*mouse models, placing an inevitable impediment to specific indirect detection of Ms monoclonal IgG Abs in the mouse brains. We provided a simple and efficient solution to the issue by the use of secondary Abs that specifically recognize Ms IgG subclasses.

## Results

### Endogenous IgG accumulation associated with Aß plaques in AD mouse models

We first assessed endogenous IgG accumulation in the brain parenchyma of an *App*-KI mouse line, *App^NL-G-F^* mouse model^30^. *App^NL-G-F^* line, which expresses humanized Aβ peptide and harbors the Swedish (KM670/671NL), Beyreuther/Iberian (I716F), and Arctic (E693G) mutations, manifests Aβ plaque deposition and cognitive impairments earlier than other *App*-KI mouse lines^30–32^. We used the mouse model aged 6 to 7 months to examine the accumulation of endogenous IgG in its brain parenchyma when memory impairment is observed^30^. The accumulation of endogenous IgG was examined by immunostaining with an anti-Ms IgG (H+L) Ab conjugated with a fluorophore (Fig. 1a). Anti-Ms IgG (H+L) Abs react with whole molecule Ms IgG which includes heavy and light chains of the immunoglobulin. Strikingly, robust immunoreactivity against Ms IgG (H+L) was found in frozen brain sections of *App^NL-G-F/NL-G-F^*mice following to the application of the anti-Ms IgG Ab (Fig. 1). Immunoreactivity against IgG, which looked like amorphous blobs, was distributed broadly in the cerebral cortex (Fig. 1a, b). Immunodetection of IgG molecules in *App^NL-G-F/NL-G-F^* mouse brains could not be explained by promiscuous binding of the Fc region of the anti-Ms IgG (H+L) Ab, because the sole application of Alexa Fluor (AF) 647 conjugated Fab fragments of an anti-Ms IgG (H+L) Ab also caused amorphous blob-like signals on frozen brain sections of *App^NL-G-F/NL-G-F^*mice (Fig. 1c). Thus, endogenous IgG in *App^NL-G-F^* model is specifically immunodetected by anti-Ms IgG (H+L) Abs via their Fab regions that recognize antigenic targets.

**Figure 1.**
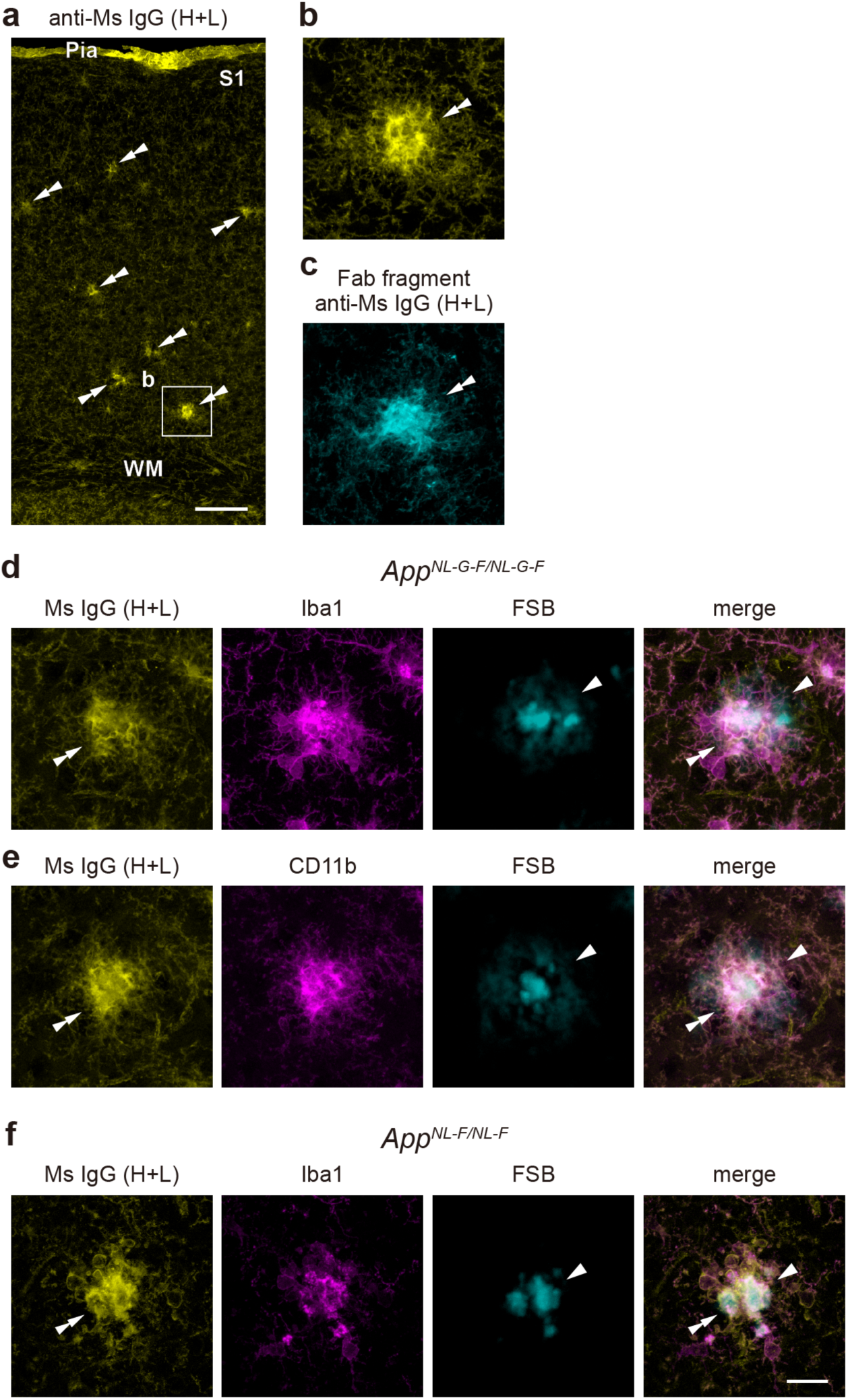
PA-IgG in *App*-KI mouse models. (**a**) Direct IF for Ms IgG (yellow) in the S1 cortex of *App^NL-G-F/NL-G-F^*mice (n = 3 mice). (**b**) Higher magnification of a rectangle in (**a**). (**c**) Ms IgG immunoreactivity in *App^NL-G-F/NL-G-F^* mice detected with AF647-counjugated Fab fragments of an anti-Ms IgG (H+L) (cyan) (n = 3 mice). (**d**-**f**) Double IF for Ms IgG (yellow) and microglial markers (Iba1: **d**, **f**, CD11b: **e**, magenta) on brain sections of *App^NL-G-F/NL-G-F^* (**d**, **e**) and *App^NL-F/NL-F^* mice (**f**) (n = 3 mice for each condition). FSB (cyan) is used for the detection of Aβ plaques. Triple merged images of Ms IgG, Iba1(**d**, **f**) or CD11b (**e**), and FSB are shown in the rightmost panels. Double arrowheads and arrowheads indicate PA-IgG and Aβ plaques, respectively. Pia: pia mater, S1: primary somatosensory cortex, and WM: white matter. Scale bars, 100 µm in (**a**) and 20 µm in (**f**).

Endogenous IgG is reportedly localized in fibrils of Aβ peptides in AD patients^33–35^. In mouse models of AD, IgG immunosignals were detected on PAM and Aβ plaques themselves^27–29^. We then sought to identify the amorphous blob-like structures labeled with anti-Ms IgG (H+L) Abs in *App^NL-G-F/NL-G-F^*mice. For this, frozen brain sections of the mouse model were exposed to Abs against Ms IgG (H+L) and ionized calcium-binding adapter molecule 1 (Iba1), a microglial marker^36^. The sections were further treated with 1-Fluoro-2,5-bis(3-carboxy-4-hydroxystyryl)benzene (FSB)^37^ for amyloid fibril labeling. In parallel with the previous studies^27,28^, fluorescent signals of an anti-Ms IgG (H+L) Ab were mostly overlapped with those of an anti-Iba1 Ab and/or FSB, indicating the accumulation of endogenous IgG on PAMs and/or Aβ plaques in *App^NL-G-F/NL-G-F^* mice (Fig. 1d). The reaction of the anti-Ms IgG (H+L) Ab with PAM and/or Aβ plaques was confirmed with another microglial marker, cluster of differentiation molecule 11b (CD11b)^38^ (Fig. 1e).

We further examined the endogenous IgG accumulation in another *App*-KI mouse line, *App^NL-F^*mouse model, which harbors the Swedish double mutations and Beyreuther/Iberian mutation in *App*^30^. A cryo-electron microscopic study shows that Aβ42 deposits in *App^NL-F^* line are identical to human type II filament fold^39^. As in *App^NL-G-F/NL-G-F^* mice, amyloid fibril labeling with FSB and immunostaining for Ms IgG (H+L) and Iba1 showed robust IgG immunoreactivity on PAM and/or Aβ plaques in the cortical parenchyma of *App^NL-F/NL-F^* mice (Fig. 1f). Taken together, these results demonstrate endogenous IgG accumulation associated with Aβ plaques in mouse models of AD.

### Hampered specific indirect detection of an Ms monoclonal IgG Ab in a mouse model of AD

PA-IgG in AD mouse models (Fig. 1) could confound specific indirect IHC of Ms monoclonal IgG Abs on brain sections of AD mouse models by causing off-target binding of secondary Abs against Ms IgG to the endogenous IgG. Indeed, when frozen brain sections of *App^NL-G-F/NL-G-F^*mice were reacted with an Ms monoclonal IgG Ab against NeuN and detected with an anti-Ms IgG (H+L) secondary Ab, the most commonly used secondary Ab for immunofluorescence (IF), amorphous blobs-like immunoreactivity was frequently found among nuclear immunoreactivity of neuronal cells (Fig. 2a, c). NeuN is a marker for nuclei of mature neurons^40^. These amorphous blob-like signals were found throughout the cortical layers of the primary somatosensory cortex (S1), in line with the broad distribution of PA-IgG in *App^NL-G-F/NL-G-F^* mouse brains (Fig. S1). In contrast, these amorphous blob-like signals were never observed in indirect IHC on *App^NL-G-F/NL-G-F^* mouse brain sections with a rabbit (Rb) polyclonal IgG Ab against NeuN (Fig. 2b, d). These amorphous blob-like signals on *App^NL-G-F/NL-G-F^* mouse brain sections are caused by off-target binding of the anti-Ms IgG (H+L) secondary Ab, because the sole application of this secondary Ab without primary Ab incubation showed amorphous blob-like immunosignals on brain sections of the AD mouse model (Fig. 2e). The sole application of the secondary Ab against Rb IgG (H+L) used in the NeuN IF did not induce such immunosignals on *App^NL-G-F/NL-G-F^* mouse brain sections (Fig. 2f). Thus, secondary Abs against Ms IgG (H+L) cause off-target binding to PA-IgG, confounding the specific indirect detection of Ms monoclonal IgG Abs in mouse models of AD. We also found immunoreactivity in the pia matter of S1 in the NeuN IF with the Ms monoclonal IgG Ab (Fig. 2a). Pial immunoreactivity was also observed in direct immunostaining for Ms IgG with the anti-Ms IgG (H+L) secondary Ab (Fig. 1a), indicating non-specific binding of the secondary Ab to the pia matter.

**Figure 2.**
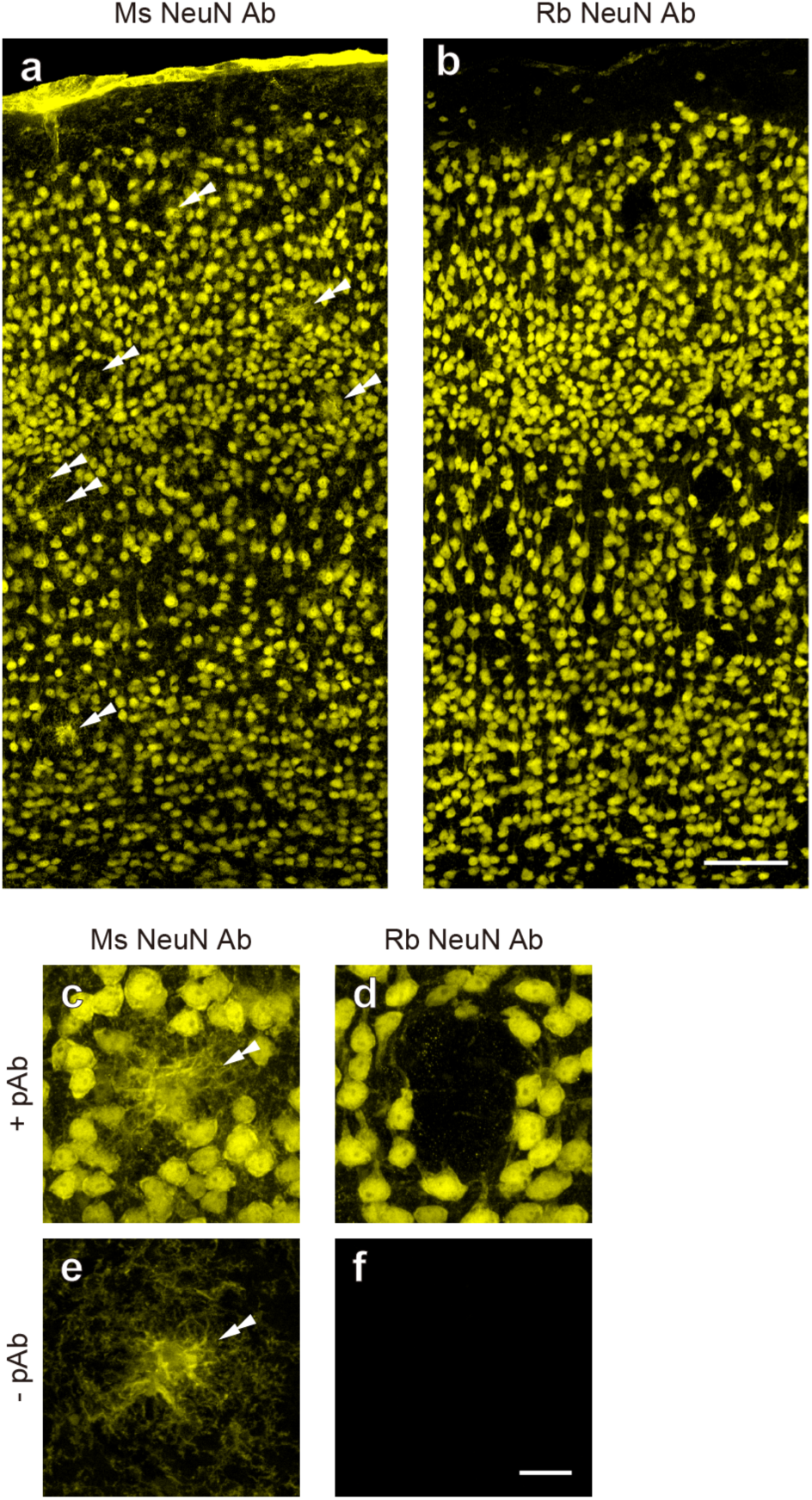
Off-target binding of secondary Abs against Ms IgG (H+L) to frozen brain sections of *App^NL-G-F/NL-G-F^* mice. (**a**, **b**) Indirect IF with Ms (**a**) and Rb (**b**) NeuN IgG Abs on brain sections of *App^NL-G-F/NL-G-F^*mice. (**c**-**f**) Higher magnifications of the indirect IF using Ms (**c**, **e**) and Rb (**d**, **f**) NeuN IgG Abs. Primary Abs are not applied in (**e**) and (**f**) (n = 3 mice for each condition). Double arrowheads indicate off-target binding of secondary Abs against Ms IgG (H+L). Scale bars, 100 µm in (**a**) and 20 µm in (**f**).

### Insufficient blocking of the off-target binding with Fab fragments of anti-Ms IgG (H+L)

Monovalent IgG Fab fragments can mask epitopes of secondary Abs to prevent secondary Abs from binding to endogenous IgG on tissue sections^41^. Especially, in indirect IHC using Ms monoclonal IgG Abs on mouse tissues, Fab fragments of IgG against Ms IgG are widely used to block off-target binding of anti-Ms IgG secondary Ab to mouse organs rich in endogenous IgG. We thus treated brain sections with excess amounts of Fab fragments of IgG against Ms IgG (H+L) (10 or 100 μg/mL) to block the off-target binding of secondary Ab against Ms IgG (H+L) in *App^NL-G-F/NL-G-F^* mice (Fig. 3). However, this blocking method with Fab fragments was not effective for the off-target binding to PA-IgG. Even after treatment with Fab fragments of anti-Ms IgG (H+L) Ab, a significant portion of the off-target binding was still detected in the *App*-KI mouse brain sections (Fig. 3). Regardless of the concentrations of the secondary Ab (1.0 or 10 μg/mL), we found amorphous blob-like signals on Aβ plaques or their vicinities in sections treated with the Fab fragments (Fig. 3a, b). Quantitative analysis demonstrated that when the secondary Ab was applied at a concentration of 10 µg/mL, the off-target binding was found on 94 ± 7.2, 84 ± 8.0, and 77 ± 6.1% of the Aβ plaques following treatment with 0, 10, and 100 μg/mL of the Fab fragment (Fig. 3a). The effects of the treatment with Fab fragments were modestly enhanced when a lower concentration of the secondary Ab (1.0 μg/mL) was applied to brain sections of *App^NL-G-F/NL-G-F^* mice (Fig. 3b). However, even in this situation, more than half of the Aβ plaques still harbored the off-target binding of the secondary Ab after pretreatment with the Fab fragments (control, 90 ± 7.0%; 10 µg/mL, 56 ± 12%; 100 μg/mL, 54 ± 5.4%). Therefore, treatment of *App^NL-G-F/NL-G-F^* mouse brain sections with Fab fragments of IgG against Ms IgG (H+L) is only partially effective for preventing the off-target binding to PA-IgG and a substantial portion of them is not affected by the conventional Ms IgG blocking method.

**Figure 3.**
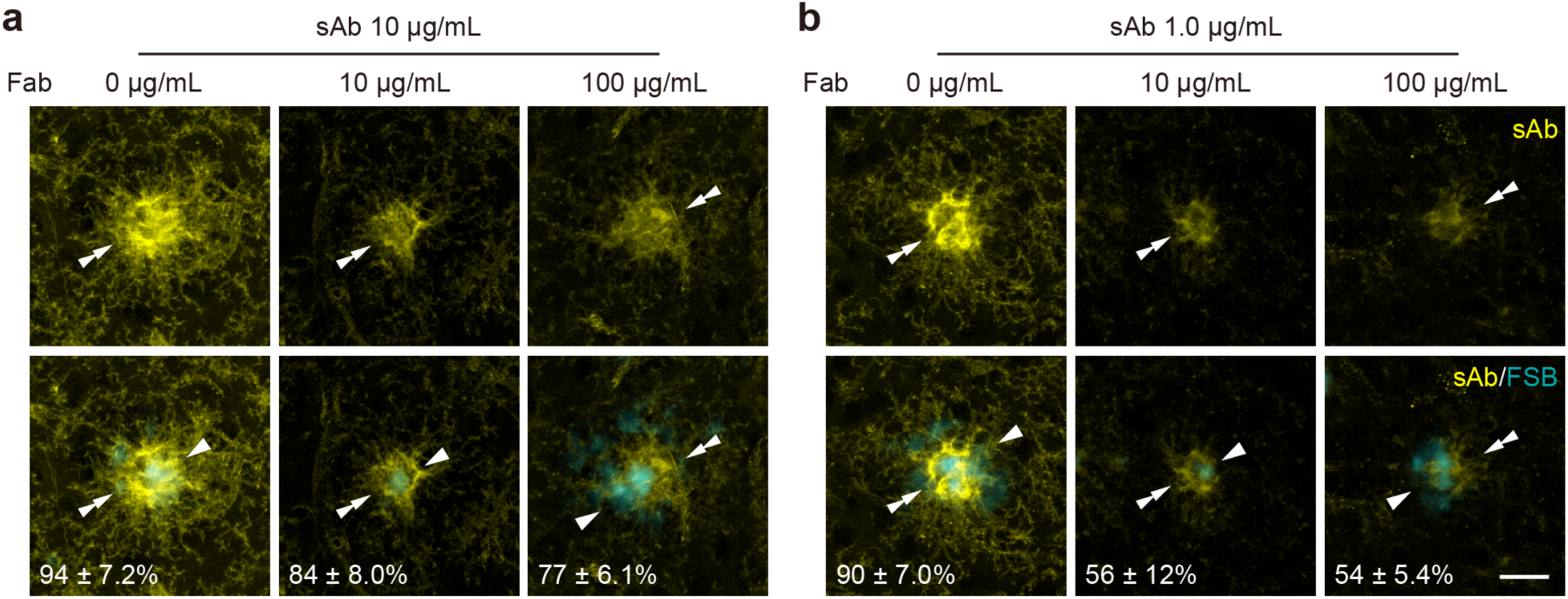
Insufficient blocking of the off-target binding by Fab fragments of anti-Ms IgG (H+L) Ab. (**a**, **b**) Applications of an AF 488 conjugated Dk anti-Ms IgG (H+L) (yellow) on *App^NL-G-F/NL-G-F^* mouse brain sections pretreated with Fab fragments of a Dk anti-Ms IgG (H+L) Ab (n = 3 mice for each condition). 10 (**a**) and 1.0 (**b**) µg/mL of the secondary Ab are applied. Aβ plaques are labeled with FSB (cyan). The top and bottom panels show images of the secondary Ab, and merged images of the secondary Ab and FSB. Concentrations of the secondary Ab and Fab fragments are indicated on the top. The percentages of Aβ plaques decorated with the secondary Ab are shown in the bottom left corner of the bottom panels. Double arrowheads and arrowheads indicate off-target binding of the secondary Ab and Aβ plaques, respectively. Scale bar, 20 µm.

### Specific detection of Ms monoclonal IgG Abs in an AD mouse model by the use of secondary Abs against Ms IgG subclasses

Nanobodies (nAbs) or variable domains of heavy chain of heavy-chain (VHH) Abs are single-domain antibody fragments derived from antigen-binding domain of heavy-chain Abs in Camelidae^42^. Owing to their superior permeability and high reproducibility^43^, accumulating studies have employed nAbs for IHC as primary and secondary Abs. We next tested whether secondary nAbs against Ms IgG caused the off-target binding to PA-IgG in *App^NL-G-F/NL-G-F^* mice. To our surprise, two of three tested secondary nAbs, nAbs against Ms IgG1 and 2b, did not show obvious off-target bindings around Aβ plaques (Fig. 4a, b). This stands in stark contrast to the aforementioned results with conventional IgG secondary Abs against Ms IgG (H+L) that react with whole molecule Ms IgG (see Fig. 2e). The absence of off-target binding was not due to their antibody forms, because amorphous blob-like signals to PA-IgG still appeared in brain sections of *App^NL-G-F/NL-G-F^* mice that were incubated with an nAb against Ms IgGκ light chain (Fig. 4c). These serendipitous findings prompted us to hypothesize that secondary Abs that specifically react with Ms IgG subclasses do not cause off-target binding to PA-IgG. To test this idea, we incubated frozen brain sections of *App^NL-G-F/NL-G-F^* mice with AF 488 conjugated whole IgG Abs against Ms IgG subclasses, including IgG1, 2a, 2b, 2c, and 3. Our hypothesis was largely correct: while an anti-Ms IgG (H+L) secondary Ab exhibited robust off-target bindings on Aβ plaques or their vicinities in the AD mouse model (Fig. 5a), secondary Abs against Ms IgG subclasses did not show amorphous blob-like bindings around them, except for a secondary Ab against Ms IgG2c (Fig. 5b–f). Amorphous blob-like signals, reflecting the off-target bindings to PA-IgG, were still prevalent in frozen brain sections of *App^NL-G-F/NL-G-F^* mice that had been treated with a secondary Ab against Ms IgG2c (Fig. 5e). This off-target binding was also observed when using another two different secondary Abs against Ms IgG2c on frozen brain sections of *App^NL-G-F/NL-G-F^* mice (Fig. S2). Based on these findings, we reasoned that the use of secondary Abs against Ms IgG subclasses, except for IgG2c, allows for specific indirect detection of Ms monoclonal IgG Abs in the AD model mouse brains.

**Figure 4.**
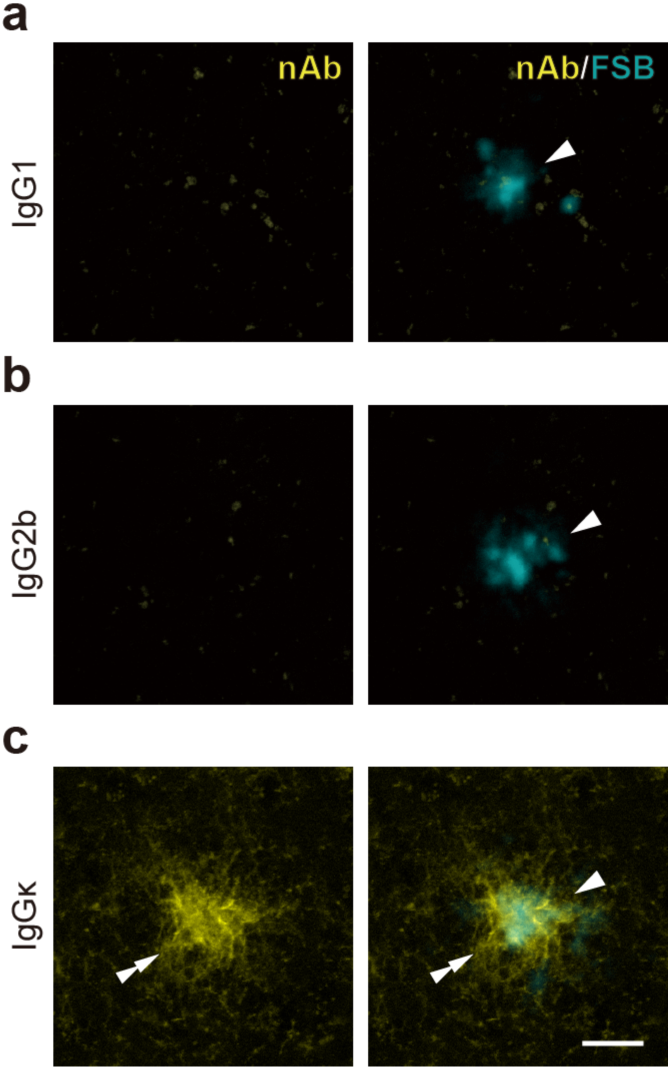
Undetectable off-target binding of secondary nAbs against Ms IgG1 and 2b to PA-IgG. (**a**-**c**) Applications of AF 488 conjugated nAbs against Ms IgG1 (**a**), 2b (**b**), and IgGκ light chain (**c**) (yellow) on frozen brain sections of *App^NL-G-F/NL-G-F^* mice (n = 3 mice for each condition). Aβ plaques are labeled with FSB (cyan). Images of secondary nAbs, and merged images of the secondary nAbs and FSB are shown in the left and right panels, respectively. Double arrowheads and arrowheads indicate off-target binding of the secondary nAb and Aβ plaques, respectively. Scale bar, 20 µm.

**Figure 5.**
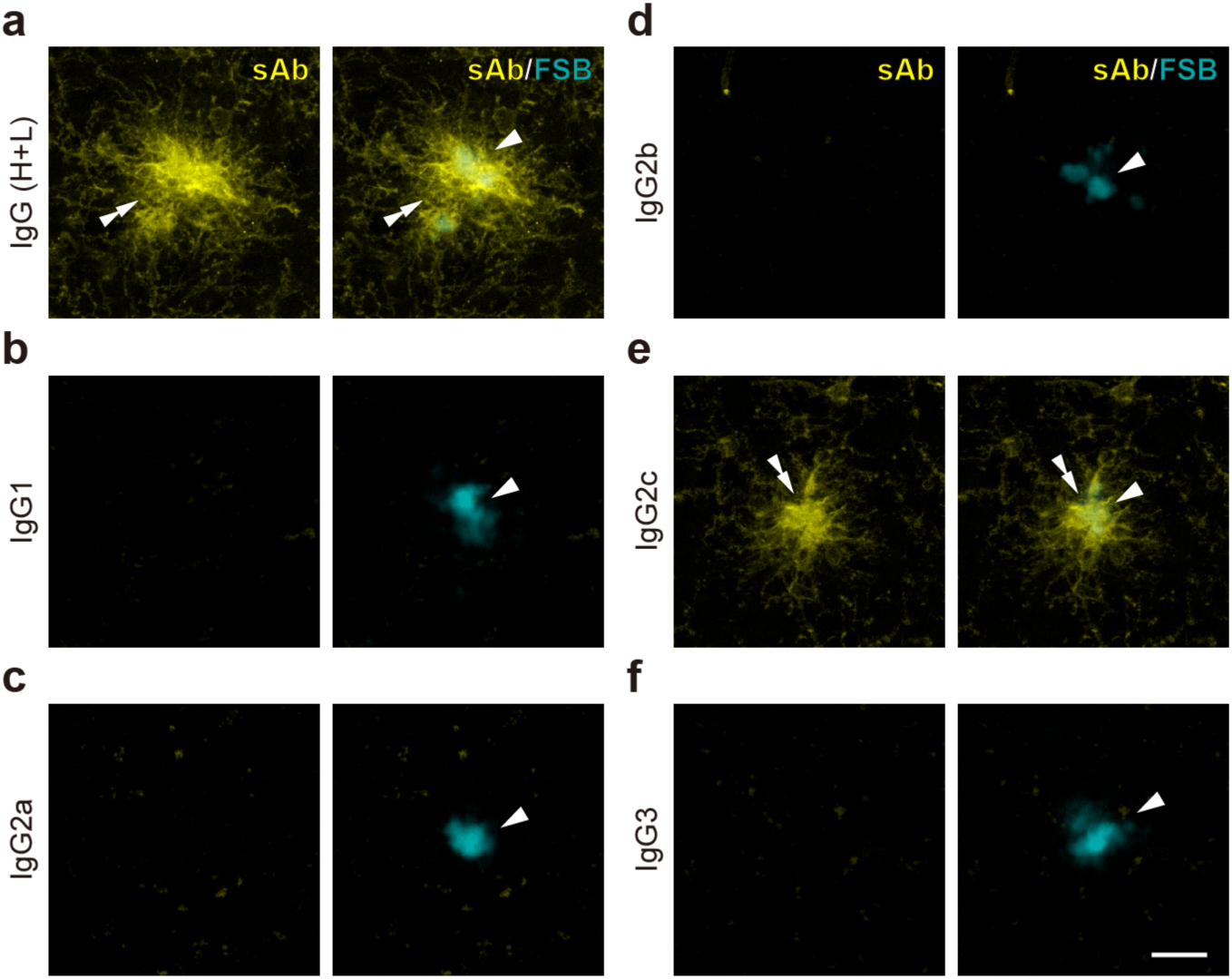
Disappearance of the off-target binding to PA-IgG by the use of secondary Abs against Ms IgG1, 2a, 2b, and 3. (**a**-**f**) Applications of secondary Abs against Ms IgG (H+L) (**a**), IgG1 (**b**), 2a (**c**), 2b (**d**), 2c (**e**) and 3 (**f**) (yellow) on brain sections of *App^NL-G-F/NL-G-F^* mice (n = 3 animals for each condition). Aβ plaques are labeled with FSB (cyan). Images of secondary Abs, and merged images of the secondary Abs and FSB are shown in the left and right panels, respectively. Double arrowheads and arrowheads indicate off-target bindings of secondary Abs and Aβ plaques, respectively. Scale bar, 20 µm.

We tested the idea by indirect IF on *App^NL-G-F/NL-G-F^*mouse brain sections using secondary Abs against Ms IgG1, 2a, 2b, and 3 (Fig. 6). We used three Ms monoclonal IgG Abs (IgG1, 2a, and 2b) against NeuN and one Ms monoclonal IgG Ab (IgG3) against microtubule-associated protein 2 (MAP2) as primary Abs in the indirect IF. MAP2 is a somatodendritic marker for neuronal cells^44^. Strikingly, secondary Abs that specifically recognized Ms IgG subclasses allowed for specific indirect detection of Ms monoclonal IgG Abs in *App^NL-G-F/NL-G-F^* mouse brains: whereas a secondary Ab against Ms IgG (H+L) caused off-target binding on Aβ plaques and their vicinities, neuronal nuclei (NeuN) and dendritic arbors (MAP2) around Aβ plaques were specifically detected without any amorphous blob-like signals by using the subclass specific secondary Abs (Fig. 6a-d). Thus, the use of secondary Abs against Ms IgG subclass enables specific detection of Ms monoclonal IgG Abs on the brain sections of an AD mouse model. Since subclass specific secondary nAbs did not cause off-target binding to PA-IgG (Fig. 4), they can be also used for specific indirect detection for Ms monoclonal IgG Abs in AD mouse model brains. However, we consider the use of whole IgG secondary Abs against Ms IgG subclasses more appropriate for specific and sensitive detection of Ms monoclonal IgG Abs in AD mouse models, because fluorescence signals generated by the whole IgG were 4.7-fold stronger than those by the nAb in NeuN IF on frozen brain sections of wild-type mice in the case of secondary Abs against Ms IgG1 (Fig. S3).

**Figure 6.**
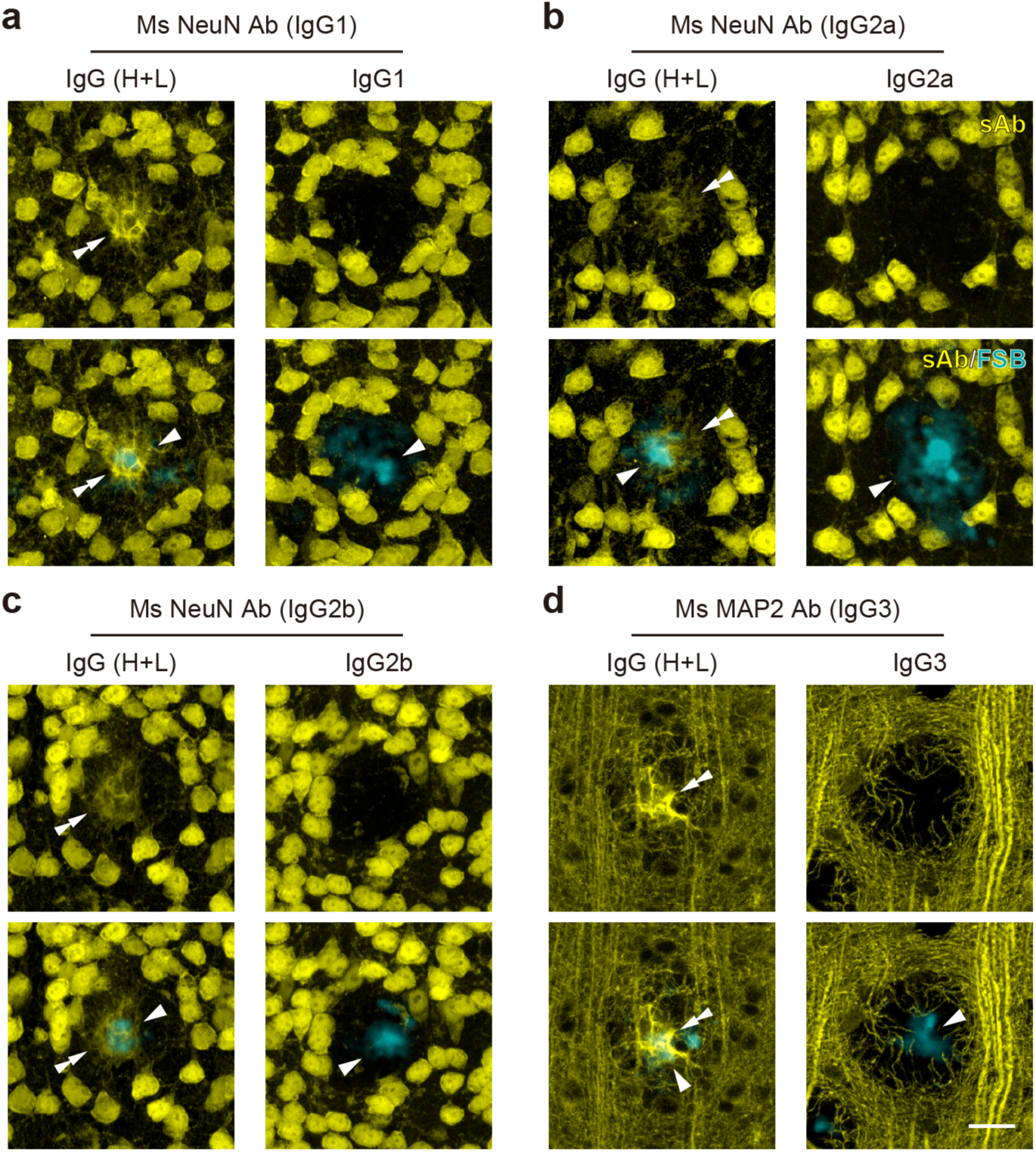
Specific detection of Ms monoclonal IgG Abs with secondary Abs against Ms IgG subclasses. (**a**-**d**) Indirect IF with Ms monoclonal IgG Abs on frozen brain sections of *App^NL-G-F/NL-G-F^* mice (n = 3 mice for each condition). Ms IgG1 (**a**), 2a (**b**), and 2b (**c**) Abs against NeuN, and an Ms IgG3 Ab against MAP2 (**d**) are used as primary Abs. Secondary Abs against Ms IgG (H+L) (left) and IgG subclasses (right) are used for their detection (yellow). Aβ plaques are labeled with FSB (cyan). The top and bottom panels show images of IF, and merged images of the secondary Abs and FSB, respectively. Double arrowheads and arrowheads indicate off-target bindings of the secondary Ab against Ms IgG (H+L) and Aβ plaques, respectively. Scale bar, 20 µm.

### Accurate immunodetection of Aβ plaque burden and phosphorylated tau accumulation by Ms IgG subclass specific secondary Abs

The off-target binding of secondary Abs against Ms IgG (H+L) to PA-IgG could cause misinterpretation on results of indirect IHC using Ms monoclonal IgG Abs in AD mouse models. Lastly, we demonstrated that outcomes of indirect IF using Ms monoclonal IgG Abs that were commonly employed in AD research were affected by secondary Abs against mouse IgG (H+L), and this was circumventable by the use of IgG subclass specific secondary Abs.

4G8, an Ms monoclonal IgG2b Ab, is a pioneering Ab for the detection of Aβ peptides that was raised against amino acid 17–24 of the Aβ peptide sequence^24,25^. Importantly, this Ab is incapable of binding to Aβ peptides of *App^NL-G-F^*model that harbors the Arctic mutation at amino acid 22 within their Aβ peptide sequence^30^. The direct method using an AF 488-conjugated 4G8 Ab confirmed these findings: while Aβ plaques in *App^NL-G-F/NL-G-F^* mice were not detected by the direct method, the 4G8 Ab clearly labeled Aβ plaques without the Arctic mutation in *App^NL-F^* model (Fig. S4). However, when a secondary Ab against Ms IgG (H+L) was employed for its indirect detection, 4G8 IF showed obvious immunosignals on Aβ plaques and their vicinities in frozen brain sections of *App^NL-G-F/NL-G-F^* mice (Fig. 7a). In contrast, indirect detection with a secondary Ab against Ms IgG2b did not show any signals in the brain of *App^NL-G-F/NL-G-F^* mice (Fig. 7a), as with the direct detection. The immunoreactivity in the indirect detection with an anti-Ms IgG (H+L) secondary Ab (Fig. 7b) was attributable to off-target binding of the secondary Ab to PA-IgG. Double indirect IF using 4G8 and an Iba1 Abs revealed that the ‘4G8 immunoreactivity’ labeled with the secondary Ab against Ms IgG (H+L) closely resembled amorphous blob-like signals caused by the sole application of secondary Abs against Ms IgG (H+L) (Fig. 2e), and this immunoreactivity mostly overlapped with that of Iba1 around Aβ plaques (Fig. 7b). Such immunoreactivity was never detected when using the Ms IgG2b subclass specific secondary Ab (Fig. 7b).

**Figure 7.**
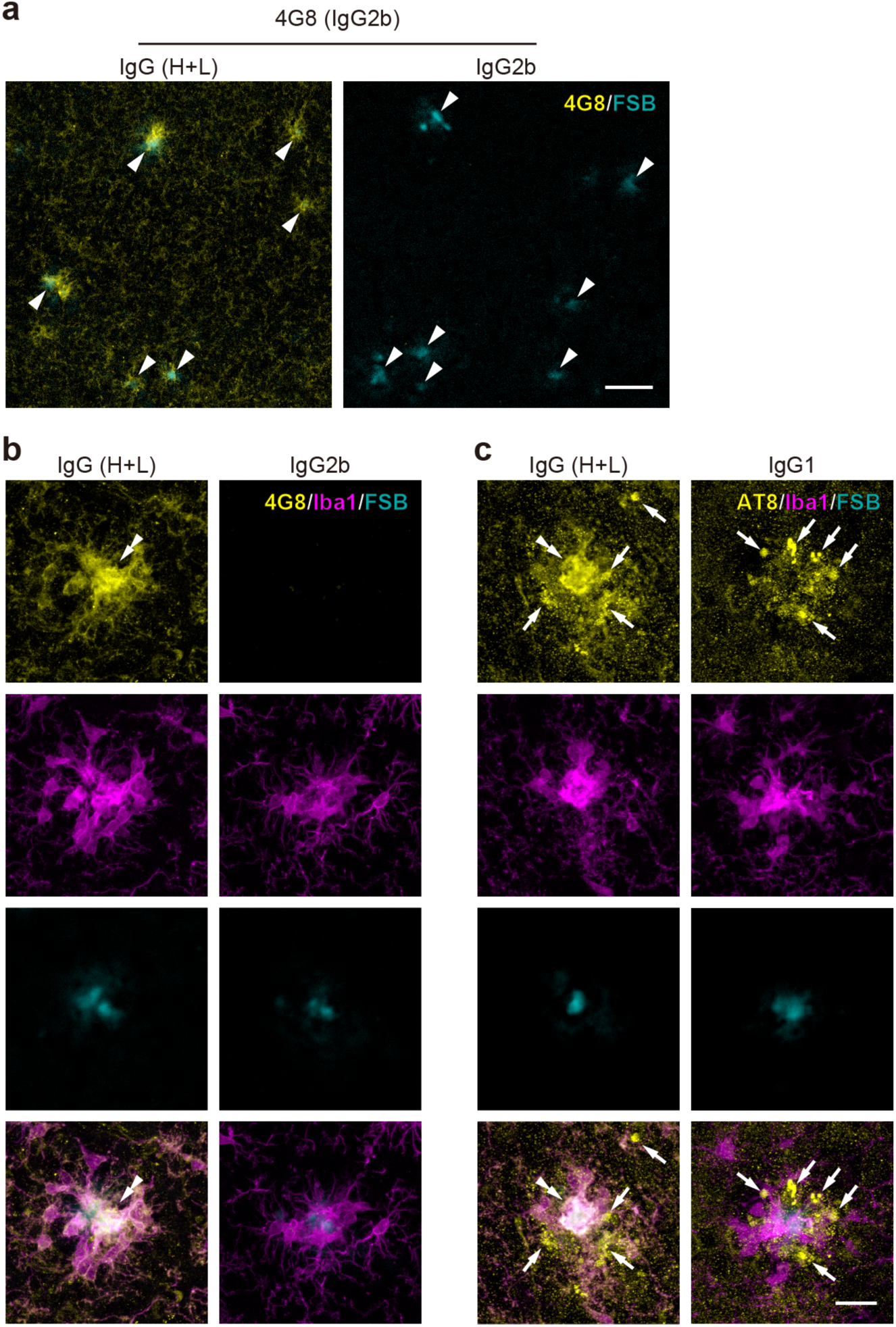
Accurate indirect detection of Ms monoclonal Abs against Aβ peptides and phosphorylated tau by the use of secondary Abs against Ms IgG subclasses. (**a**) Indirect IF of 4G8 Ab (yellow) using secondary Abs against Ms IgG (H+L) (left) and IgG2b (right) on frozen brain sections of *App^NL-G-F/NL-G-F^*mice (n = 3 mice for each condition). Aβ plaques are labeled with FSB and indicated by arrowheads (cyan). (**b**) Triple labeling of 4G8 Ab (yellow), an Iba1 Ab (magenta), and FSB (cyan) on frozen brain sections of *App^NL-G-F/NL-G-F^* mice (n = 3 mice for each condition). AF 488 conjugated anti-Ms IgG (H+L) (left) and Ms IgG2b Abs (right) are used as secondary Abs. Triple merged images of 4G8 Ab, the Iba1 Ab, and FSB are shown in the bottommost panel. Double arrowheads indicate off-target binding to PA-IgG. (**c**) Triple labeling of AT8 Ab (yellow), the Iba1 Ab (magenta), and FSB (cyan) on frozen brain sections of *App^NL-G-F/NL-G-F^* mice (n = 3 mice for each condition). AT8 Ab is indirectly detected with secondary Abs against IgG (H+L) (left) and IgG1 (right). The bottommost panel show triple merged images of AT8 Ab, the Iba1 Ab, and FSB. Double arrowheads and arrows show off-target binding to PA-IgG, and immunosignals in dystrophic neurites, respectively. Scale bars, 50 µm in (**a**) and 20 µm in (**c**).

NFTs, which are caused by intracellular accumulation of phosphorylated and misfolded tau proteins, represent a major histopathological hallmark of AD^4,45^. AT8 Ab with Ms IgG1 subclass is the first Ab that specifically recognizes phosphatase-sensitive epitopes of tau proteins^26^. AT8 Ab has been a golden standard for immunodetection of pathological tau proteins. In AD mouse models with β-amyloidosis in their brains, such as *App^NL-G-F/NL-G-F^* model, phosphorylated tau proteins typically accumulate in dystrophic neurites^46^ around Aβ plaques without NFT formation^47,48^. However, indirect detection of AT8 with a secondary Ab against Ms IgG (H+L) showed amorphous blob-like signals near Aβ plaques, in addition to immunosignals in dystrophic neurites (Fig. 7c). Strikingly, the amorphous blob-like AT8 signals were largely localized to the region labeled with an Iba Ab at the immediate vicinity of Aβ plaques (Fig. 7c). In contrast, AT8 immunoreactivity was mostly restricted to dystrophic neurites near Aβ plaques, when a secondary Ab against Ms IgG1 was used for its detection (Fig. 7c). Notably, the AT8 immunoreactivity was rarely found in regions where immunoreactivity for Iba1 was detected (Fig. 7c), indicating that the ‘AT8 immunoreactivity’ on PAMs in the indirect IHC with a secondary Ab against Ms IgG (H+L) was likely due to off-target binding to PA-IgG in *App^NL-G-F/NL-G-F^* mice. Thus, indirect detection with secondary Abs against Ms IgG (H+L) could lead to faulty conclusions regarding Aβ plaque burden and phosphorylated tau accumulation in AD mouse models, and the use of secondary Abs that specifically recognize Ms IgG subclasses circumvents the inevitable misinterpretations caused by off-target binding to PA-IgG in AD mouse models exhibiting β-amyloidosis.

## Discussion

In the present study, we report endogenous IgG accumulation associated with Aβ plaques and its impact on indirect IHC with Ms monoclonal IgG Ab in AD mouse models. Endogenous IgG was accumulated on PAM and/or Aβ plaques in AD mouse models, which placed an inevitable impediment to the specific indirect detection of Ms monoclonal IgG Ab on frozen brain sections of *App^NL-G-F/NL-G-F^* mice (Fig. 1, 2). Preincubation with Fab fragments of anti-Ms IgG (H+L) Abs was insufficient for blocking the off-target binding of a secondary Ab against Ms IgG (H+L) (Fig. 3). Strikingly, we found that nAbs and whole IgG Abs against Ms IgG subclasses, with the exception for IgG2c, did not cause the off-target binding (Fig. 4, 5), and these whole IgG secondary Abs allowed for specific indirect detection of Ms monoclonal IgG Abs on *App^NL-G-F/NL-G-F^*mouse brain sections (Fig. 6, 7). We further demonstrated that indirect detection with a conventional secondary Ab against Ms IgG (H+L) Ab could lead to faulty conclusions regarding Aβ plaque burden and phosphorylated tau accumulation in AD model mice, which can be avoided by using IgG subclass specific secondary Abs (Fig. 7).

Our IHC analysis using fluorophore-conjugated Abs against Ms IgG (H+L) Ab showed that endogenous IgG accumulated on PAM and/or Aβ plaques in *App^NL-F^*and *App^NL-G-F^* mouse models (Fig. 1). Endogenous IgG accumulation within Aβ plaques has been reported in patients with AD^33–35^. Importantly, PA-IgG is not exclusive to these two *App*-KI mouse models. Accumulation of endogenous IgG is reported in brain sections of at least another two mouse models of AD, 5xFAD^27,28^ and arcAβ^29^, suggesting that endogenous IgG accumulation within and around Aβ plaques is a general phenomenon associated with β-amyloidosis. PA-IgG could hamper specific indirect detection with Ms monoclonal IgG Abs in the brain of AD mouse models by causing off-target binding of anti-Ms IgG secondary Abs that react with endogenous Ms IgG. However, numerous studies have shown specific indirect detection of Ms monoclonal IgG Abs in AD model mouse brains even where the accumulation of endogenous IgG is found on PAM and/or Aβ plaques^30,49,50^. Notably, the off-target binding of secondary Abs against Ms IgG (H+L) is prevalent on frozen brain sections^27–29^. Frozen sections more often retain cellular and molecular integrity compared to paraffin sections. All of studies that report the accumulation of endogenous IgG used frozen brain sections prepared from the AD mouse models^27–29^. The off-target binding of anti-Ms IgG (H+L) Abs might be less frequent on paraffin sections, allowing for specific indirect detection by the use of secondary Abs against Ms IgG (H+L). Indeed, Whyte et al. show specific detection of an Ms monoclonal IgG Ab against NeuN (MAB377, Millipore), which is also employed in this study, on paraffinized brain sections of *App^NL-G-F/NL-G-F^* mice using an anti-Ms IgG (H+L) secondary Ab^51^. Specific indirect detection on paraffin sections of *App^NL-G-F/NL-G-F^*mice using a secondary Ab against Ms IgG (H+L) has also been reported in IF for Parvalbumin, a marker for a subpopulation of cortical and hippocampal inhibitory interneurons^52^. IHC procedures on paraffin sections, such as dehydration, paraffin wax embedding, dewaxing, rehydration, and antigen retrieval, might disturb the antigen-antibody interaction between endogenous Ms IgG and secondary Abs against Ms IgG (H+L) that is preserved in frozen brain sections of AD mouse models. The appearance of the off-target binding is also likely to be dependent on primary Abs used in IHC. Several studies have shown specific labeling of Ms monoclonal IgG Abs using anti-Ms IgG (H+L) secondary Abs without obvious off-target binding on frozen brain sections of *App^NL-G-F/NL-G-F^* mice^53–55^.

We found that secondary Abs against Ms IgG1, 2a, 2b, and 3 did not cause obvious off-target binding on *App^NL-G-F/NL-G-F^* mouse brain sections (Fig. 4, 5). Although the mechanisms underlying the disappearance of the off-target binding by the use of the secondary Abs have not totally elucidated, we propose the following potential mechanisms which can be responsible for the vanishment. For IgG2a, lack of the IgG2a gene in the C57BL/6J mouse^56^, which is the background strain of *App^NL-G-F/NL-G-F^*mice used in this study, can explain the disappearance of the off-target binding. Actually, we confirmed the lack of the IgG2a gene from *App^NL-G-F/NL-G-F^* mice used in this study by polymerase chain reaction (PCR) genotyping with primers targeting the IgG2a gene (data not shown). Two possible mechanisms can be considered for IgG1, 2b, and 3. The first is that IgG1, 2b, and 3 are not accumulated on PAM and/or Aβ plaques in the AD mouse model. Given the influx of exogenously administrated IgG1 and 2b Abs into the mouse brain parenchyma^57,58^, endogenous IgG1 and 2b might flow into the brain parenchyma, but fail to accumulate on PAM and/or Aβ plaques in the *App*-KI mice. The Fc gamma receptor (FcγR), which can bind to extracellular IgG molecules, has four family members in mice^59,60^. The binding affinity of IgG to FcγRs varies among IgG subclasses^61^. Of the four FcγRs, FcγRI and IV are classified into high-affinity receptors, and show higher affinities for IgG2a and 2c than other subclasses^62–64^. Thus, IgG1 and 2b that flowed into the brain parenchyma might not fully bind to the surface of PAM and/or Aβ plaques of *App^NL-G-F/NL-G-F^* mice. It remains uncertain whether IgG3 can flow into the brain parenchyma. Additionally, IgG3 has the lowest affinity for all FcγRs among Ms IgG subclasses^61,64^. The second is that IgG1, 2b, and 3 accumulated on PAM and/or Aβ plaques cannot be detected by the secondary Abs against Ms IgG subclass. In this scenario, IgG1, 2b, and 3 flow into the brain parenchyma and accumulate within and/or around Aβ plaques, but the subclass-specific secondary Abs used in this study failed to bind the accumulated IgG molecules. The FcγR-IgG binding site locates close to the IgG hinge region where Ms IgG subclasses exhibit the most conspicuous differences in their amino acid sequences^65^. Thus, the epitopes of the subclass-specific secondary Abs might be masked by the interactions between IgGs and FcγRs, preventing these subclass-specific secondary Abs from binding to PA-IgG in AD mouse models.

Although other strategies, such as the direct method, secondary nAbs against Ms IgG subclasses, and Fabulights, can be employed for specifically detecting Ms monoclonal IgG in AD mouse models, we think indirect detection using whole IgG secondary Abs against Ms IgG subclasses more appropriate for specific and sensitive detection of Ms monoclonal IgG Abs. The direct method, which uses primary Abs directly conjugated with fluorophores or haptens, labels target antigens without the use of secondary Abs that can cause off-target binding to PA-IgG. However, it has lower sensitivity than the indirect method^17,18^. Furthermore, the direct method requires labeling of primary Abs with fluorophores or haptens, which is costly, laborious, and time consuming. Secondary nAbs and Fabulights that specifically recognize Ms IgG subclass can also be used for specific detection of Ms monoclonal IgG Abs in AD mouse models. Indeed, secondary nAbs against IgG1 and 2b did not show obvious off-target binding to frozen brain sections of *App^NL-G-F/NL-G-F^*mice (Fig. 4a, b). Fabulights are fluorophore or hapten-conjugated Fab fragments of secondary IgG Abs that specifically bind the Fc region of IgG molecules (https://www.jacksonimmuno.com/technical/products/groups/fab/fabulight). Ms monoclonal IgG Abs can be labeled with Fabulights prior to reaction with tissue sections, avoiding the application of unbound anti-Ms IgG secondary Abs that can react with PA-IgG. However, the signal intensity of indirect detection using a secondary nAb and Fabulight was significantly lower than that using a whole IgG secondary Ab (Fig. S3). Additionally, IHC procedures using whole IgG secondary Abs against Ms IgG subclasses are the same as conventional indirect IHC without additional labeling of primary Abs, and are easily scalable to multiplex labeling and other histochemical techniques.

The appearance of amorphous blob-like signals on *App^NL-G-F/NL-G-F^*mouse brain sections treated with a secondary Ab against Ms IgG2c indicates the off-target binding of the Ab to PA-IgG (Fig. 5e). This cannot be explained by variations in anti-Ms IgG2c Abs, because another two different secondary Abs against Ms IgG2c showed off-target binding to frozen brain sections of *App^NL-G-F/NL-G-F^*mice (Fig. S2). Moreover, the off target-binding hampered specific detection of an Ms monoclonal IgG2c Ab on frozen brain sections of the *App*-KI mouse model (Fig. S5). We consider that the direct method or detection with Fabulights should be employed in indirect IHC using Ms monoclonal IgG2c Abs on frozen brain sections of AD mouse models. Enhancing the specificity of anti-Ms IgG2c Abs might prevent the off-target binding and enable specific detection of Ms monoclonal IgG2c Ab in AD mouse models while maintaining its high sensitivity.

Indirect detection using secondary Abs against Ms IgG (H+L) can lead to erroneous conclusions regarding IHC outcomes in AD mouse models. Indeed, we showed that the use of a secondary Abs against Ms IgG (H+L) can result in fatal misinterpretations regarding Aβ plaque burden and phosphorylated tau accumulation in *App^NL-G-F/NL-G-F^* mouse brains (Fig. 7), expressing serious concerns about the validity of indirect IHC using Ms monoclonal IgG Ab in AD mouse models. We further demonstrated that these misinterpretations can be avoided by the use of secondary Abs against Ms IgG subclass (Fig. 7). Secondary Abs that specifically recognize Ms IgG subclass should be employed for indirect detection of Ms monoclonal IgG Abs in other AD mouse models. Our method using secondary Abs against Ms IgG subclasses might be effective in the cases where the accumulation of endogenous Ms IgG hampers the specific detection of Ms monoclonal IgG Abs. The use of these secondary Abs against Ms IgG subclasses would accelerate AD research by expanding the choice of Abs available in experimental studies of the diseases.

## Methods

### Animals

All animal experiments were approved by the Institutional Animal Care and Use Committees of Juntendo University (Approval No. 2023194) and Animal Experiment Committee of RIKEN (Approval No. 2023-2-017-3), and performed in accordance with Fundamental Guidelines for Proper Conduct of Animal Experiments by the Science Council of Japan (2006). All animal procedures were conducted in compliance with ARRIVE (Animal Research: Reporting In Vivo Experiments) guidelines.

Male and female 9-week-old C57BL/6J mice (Nihon SLC), male 14-month-old *App^NL-F/NL-F^* mice^30^ (RBRC06343, RIKEN BioResource Center), and male and female 6 to 7- or 12-month-old *App^NL-G-F/NL-G-F^* mice^30^ (RBRC06344, RIKEN BioResource Center) were used. *App^NL-F^* and *App^NL-G-F^* lines were maintained on the C57BL/6J background. Mice were housed under a 12/12 h light/dark cycle with access to food and water *ad libitum*. The genotype of *App^NL-F/NL-F^* and *App^NL-G-F/NL-G-F^*mice was determined by PCR of tail DNA with primers listed in Supplementary Table 1.

### Tissue preparation

Mice were deeply anesthetized by an intraperitoneal injection of an overdose of sodium pentobarbital (200 mg/kg; Somnopentyl, Kyoritsu Seiyaku). The depth of anesthesia was confirmed by the absence of eye-blink reflexes and toe-pinch withdrawal. The right atrial appendage was cut out, and the mice were transcardially perfused with 20 mL of ice-cold phosphate buffered saline (PBS), followed by the same volume of ice-cold 4% paraformaldehyde (PFA) (1.04005.1000, Merck Millipore) in 0.1 M phosphate buffer (PB; pH 7.4), through the left ventricle. Brains of the mice were removed, postfixed in the same fixative for 24 h at 4°C, and cryoprotected in 30% (w/v) sucrose in 0.1 M PB at 4 °C. Then, the brains were separated into the two hemispheres, and cut coronally at 30-µm intervals on a freezing microtome (REM-710; Yamato Kohki Industrial), as described previously^66^. The sections were stored in 0.2% sodium azide in PBS at 4°C until use.

### Immunofluorescence

Immunofluorescence was performed by a free-floating method at 20–25°C. Coronal brain sections at the level of the hippocampus were subjected to IF. The primary and secondary Abs used in this study are listed in Tables 1 and 2, respectively.

**Table 1:** Primary Abs used in this study.

| Host | Clonality | Antigen | IgG subclass | Manufacturer, Cat. No. | RRID | Concentration /dilution |
| --- | --- | --- | --- | --- | --- | --- |
| Ms | Monoclonal | Aβ | IgG2b | BioLegend, 800708 | AB_2734547 | 1:1000 |
| Ms | Monoclonal | Aβ | IgG2b | BioLegend, 800714 | AB_2721287 | 5.0 μg/mL |
| Ms | Monoclonal | MAP2 | IgG3 | Genetex, GTX634473 | AB_2888458 | 1.2 μg/mL |
| Ms | Monoclonal | MeCP2 | IgG2c | abcam, ab252840 | N/A<sup>1</sup> | 0.96 μg/mL |
| Ms | Monoclonal | NeuN | IgG1 | Merck Millipore, MAB377 | AB_2298772 | 1.0 μg/mL |
| Ms | Monoclonal | NeuN | IgG2a | Cell Signaling Technology, 94403S | AB_2904530 | 1:600 |
| Ms | Monoclonal | NeuN | IgG2b | Thermo Fisher Scientific, MA533103 | AB_2802653 | 1.0 μg/mL |
| Ms | Monoclonal | Phosho-Tau | IgG1 | Thermo Fisher Scientific, MN1020 | AB_223647 | 0.20 μg/mL |
| Rb | Polyclonal | Iba1 | N/A<sup>1</sup> | FUJIFILM Wako Pure Chemical Corporation, 019-19741 | AB_839504 | 1:1000 |
| Rb | Polyclonal | NeuN | N/A<sup>1</sup> | Merck Millipore, ABN78 | AB_10807945 | 1.0 μg/mL |
| Rt | Monoclonal | CD11b | IgG2b | Thermo Fisher Scientific, 14-0112-82 | AB_467108 | 1.0 μg/mL |

<sup>1</sup>N/A: not available.

**Table 2:** Secondary Abs used in this study.

| Host | Antigen | Type of antibody | Conjugate | Manufacturer, Cat. No. | RRID | Concentration |
| --- | --- | --- | --- | --- | --- | --- |
| Al<sup>1</sup> | Ms IgG1 Fc region | Nanobody | AF 488 | Thermo Fisher Scientific, SA5-10329 | AB_2868376 | 1.0 µg/mL |
| Al<sup>1</sup> | Ms IgG2b Fc region | Nanobody | AF 488 | Thermo Fisher Scientific, SA5-10335 | AB_2868382 | 0.50 µg/mL |
| Al<sup>1</sup> | Ms IgGκ chain | Nanobody | AF 647 | Synaptic Systems, N1202-AF647-S | N/A<sup>2</sup> | 10 nM |
| Dk | Ms IgG (H+L) | Fab fragment | AF 647 | Jackson ImmunoResearch, 715-607-003 | AB_2340867 | 8.5 µg/mL |
| Dk | Ms IgG (H+L) | Whole IgG | AF 488 | Thermo Fisher Scientific, A21202 | AB_141607 | 1.0 or 10 µg/mL |
| Dk | Rb IgG (H+L) | Whole IgG | AF 488 | Thermo Fisher Scientific, A21206 | AB_2535792 | 10 µg/mL |
| Dk | Rb IgG (H+L) | Whole IgG | AF 568 | Thermo Fisher Scientific, A10042 | AB_2534017 | 10 µg/mL |
| Dk | Rt IgG (H+L) | Whole IgG | Rhodamine Red-X | Jackson ImmunoResearch, 712-295-153 | AB_2340676 | 15 µg/mL |
| Gt | Ms IgG1 | Whole IgG | AF 488 | Thermo Fisher Scientific, A21121 | AB_2535764 | 1.0 µg/mL |
| Gt | Ms IgG1 Fc region | Fab fragment (Fabulight) | AF 488 | Jackson ImmunoResearch, 115-547-185 | AB_2632534 | 1.0 µg/mL |
| Gt | Ms<br>IgG2a Fc<br>region | Whole IgG | AF 488 | Jackson<br>ImmunoResearch,<br>115-545-206 | AB_2338855 | 2.3 µg/mL |
| Gt | Ms<br>IgG2b | Whole IgG | AF 488 | Thermo Fisher<br>Scientific,<br>A21141 | AB_141626 | 1.0 µg/mL |
| Gt | Ms<br>IgG2c | Whole IgG | Biotin | Bethyl<br>Laboratories,<br>A90-136B | AB_10634640 | 4.0 µg/mL |
| Gt | Ms<br>IgG2c Fc<br>region | Whole IgG | AF 488 | Jackson<br>ImmunoResearch,<br>115-545-208 | AB_2338857 | 1.5 µg/mL |
| Gt | Ms<br>IgG2c<br>heavy<br>chain | Whole IgG | HRP | abcam, ab97255 | AB_10680258 | 4.0 µg/mL |
| Gt | Ms IgG3 | Whole IgG | AF 488 | Thermo Fisher<br>Scientific,<br>A21151 | AB_2535784 | 1.0 µg/mL |
| Gt | Ms IgG<br>(H+L) | Whole IgG | AF 488 | Thermo Fisher<br>Scientific,<br>A11001 | AB_2534069 | 1.0 µg/mL |
| Gt | Ms IgG<br>(H+L) | Whole IgG | AF 568 | Thermo Fisher<br>Scientific,<br>A11031 | AB_144696 | 10 µg/mL |

<sup>1</sup>Al, Alpaca
<sup>2</sup>N/A: not available.

For detection of PA-IgG in AD mouse models, brain sections were washed twice in 0.3% Triton X-100 in PBS (PBS-X) for 10 min, blocked in PBS-X containing 10% normal donkey (Dk) serum (S30-100ML, Merck Millipore) (NDkS) or normal goat (Gt) serum (S26-100ML, Merck Millipore) (NGtS) for 1 h, and reacted with anti-Ms IgG (H+L) Abs in PBS-X containing 0.12% λ-carrageenan (22049, Sigma-Aldrich) and 1% NDkS (PBS-XCD) or NGtS (PBS-XCG) for 4 h. The sections were washed twice in PBS-X for 10 min, mounted onto aminopropyltriethoxysilane (APS)-coated glass slides (APS-01, Matsunami Glass), and coverslipped with Fluoromount/PlusTM (K048, Diagnostic BioSystems).

For direct IF with 4G8 Ab, brain sections were immunostained as described above, except for overnight incubation with the primary Ab. For indirect IF, following to washes and blocking as above, brain sections were reacted overnight with primary Abs in PBS-XCD or - XCG. The sections were washed twice in PBS-X for 10 min, and reacted with secondary Abs in PBS-XCD or -XCG for 4 h. Following to IF, the sections were washed twice in PBS-X for 10 min, and treated with FSB (10 μg/mL, 341-90811, Dojindo) in PBS-X for 30 min. The sections were washed, mounted, and coverslipped as described above.

To evaluate the off-target binding of secondary Abs, brain sections were incubated in PBS-XCD or -XCG overnight following to blocking in PBS containing 10% NDkS or NGtS for 1 h. The sections were washed twice in PBS-X for 10 min, and reacted with secondary Abs in PBS-XCD or -XCG for 4 h. For detection of a biotinylated secondary Ab, the sections were reacted with AF 488 conjugated streptavidin (0.25 μg/mL, S11223, Thermo Fisher Scientific) in PBS-XCD for 4 h. For color development of a horse radish peroxidase (HRP)-conjugated secondary Ab, Fluorochromized Tyramide-Glucose Oxidase (FT-GO) signal amplification method was carried out with CF488A tyramide as described previously^18^. After color development, the brain sections were washed, labelled with FSB, mounted, and coverslipped as described above.

For detection with the Fabulight Ab, following to an incubation of the primary and Fabulight Abs at an equal weight ratio, the resulting complex of the primary Ab with the Fabulight Ab was applied to brain sections containing the S1.

### Blocking with Fab fragments of an anti-Ms IgG Ab

Fab fragments of a Dk anti-Ms IgG (H+L) Ab (715-007-003, Jackson ImmunoResearch Laboratories) were used. Brain sections of *App^NL-G-F/NL-G-F^* mice were sequentially blocked in PBS-X containing 0, 10, or 100 µg/mL of the Fab fragments of a Dk anti-Ms IgG Ab and 10% NDkS in PBS-X for 1 h each. The sections were incubated in PBS-XCD overnight, and incubated with 1.0 or 10 µg/mL of an AF 488 conjugated Dk anti-Ms IgG (H+L) Ab (A21202, Thermo Fisher Scientific) for 4 h. They were labelled with FSB, mounted, and coverslipped as described above.

### Image acquisition and processing

Images of brain sections were acquired with a confocal laser scanning microscope (TCS SP8, Leica Microsystem) using a 16× multi-immersion objective (HC FLUOTAR 16×/0.60 IMM CORR VISIR, NA = 0.60, Leica Microsystems) and a 25× water-immersion objective (HC FLUOTAR L 25×/0.95 W VISIR, NA = 0.95, Leica Microsystems) lenses. Images with a 1024 × 1024 pixel resolution were obtained with a zoom factor of 1.0 or 3.0 using a photon-counting mode of a hybrid detector (Leica Microsystems). The confocal z-step sizes were set to 1.5 and 4.0 µm for the 25× and 16× objective lenses. The confocal pinholes were set to 1.0 for IF and 4.0 for FSB labeling. FSB, AF 488, 568, and 647 were excited with 405-, 488-, 552-, and 638-nm lasers, and their emission prism windows were set to 410-510, 495-560, 577-642, and 645-710 nm, respectively. Acquired images were tiled, stitched, and maximum intensity projections (MIP) images were created using Leica Application Suite X (LAS X) software (ver. 3.7.5.24914, Leica Microsystems). Fiji (ImageJ) software^67^ (ver. 1.54f, National Institutes of Health) was used to crop images and create MIP images of three optical sections immunolabeled for MAP2 (3.0 μm). The brightness and contrast of images were adjusted globally using Adobe Photoshop (ver. 24.7.0, Adobe).

### Quantitative analysis

NeuN IF in the S1 were used to assess off-targeting binding of a Dk anti-Ms IgG (H+L) secondary Ab (A21202, Thermo Fisher Scientific) and a Dk anti-Rb IgG (H+L) secondary Ab (A21206, Thermo Fisher Scientific). The off-targeting binding of the secondary Abs appeared as some kind of amorphous blobs on brain sections of *App^NL-G-F/NL-G-F^*mice (Fig. 2, S1). The number of off-targeting binding sites was counted by visual inspection. The region of interest (ROI) in the S1 was defined as follows: first, a line with a length of 1,500 μm was drawn tangentially at the pial surface of S1. Then, two lines were drawn perpendicularly to the tangential line at the both ends of the tangential line. Finally, the lower edge of the white matter was traced between the two perpendicular lines. The region enclosed by the four lines was selected as the ROI (Fig. S1).

To assess the effects of blocking by pretreatment with Fab fragments of Dk anti-Ms IgG (H+L) Ab, Aβ plaques were randomly selected from the S1 (10 to 17 plaques in each brain section). The number of Aβ plaques decorated with an AF 488 Dk anti-Ms IgG (H+L) Ab (A21202, Thermo Fisher Scientific) among total Aβ plaques was counted, and the proportion of these Aβ plaques in the total number of Aβ plaques was calculated. Visual inspection was used to judge the presence or absence of signals.

The fluorescence intensity (arbitrary units [AU]) of NeuN IF using anti-Ms IgG1 secondary Abs with different Ab forms was measured, as described previously^18^. Optical section images at 3 μm depth from the surface were used for quantification. Fluorescence intensity of NeuN positive cells with visible nucleoli in layer 2/3 of the S1 was measured using the ImageJ software. Nucleoli were visualized using NeuroTrace 530/615 Red Fluorescent Nissl Stain (N21482, Thermo Fisher Scientific).

### Statistical analyses

Statistical analyses were performed using GraphPad Prism 7 software (ver. 7.05.237, GraphPad Software). One-way ANOVA followed by Tukey’s post hoc test was used to compare independent groups. All tests were two-sided, and *P* < 0.05 was considered statistically significant.

## Data availability

The datasets generated and/or analyzed during the current study and all biological materials described in this article are available from the corresponding author on reasonable request.

## Acknowledgements

The authors thank Drs. Takashi Saito (Nagoya City University, Japan) and Takaomi C Saido (RIKEN Brain Science Institute, Japan) for sharing *App^NL-F^* and *App^NL-G-F^* mice, Masakazu Miyajima (Juntendo University, Japan) and Madoka Nakajima (Juntendo University, Japan) for breeding and maintaining *App^NL-G-F^* mice, and Editage (www.editage.com) for English language editing. This work was supported by JSPS KAKENHI (JP22K20692 to S.I.; 23K06310 to K.Y.; JP20K07743 to M.K.; JP21H02592 to H. Hioki) from the Japan Society for the Promotion of Science (JSPS). This work was also supported by Brain Mapping by Integrated Neurotechnologies for Disease Studies (Brain/MINDS) from the Japan Agency for Medical Research and Development (AMED; JP15dm0207001 to A.M.; JP21dm0207112 to H. Hioki), Moonshot R&D from the Japan Science and Technology Agency (JST; JPMJMS2024 to H. Hioki), and Fusion Oriented Research for disruptive Science and Technology (FOREST) from JST (JPMJFR204D to H. Hioki).

## Author information

### Authors and Affiliations

**Department of Neuroanatomy, Juntendo University Graduate School of Medicine, Bunkyo-Ku, Tokyo 113-8421, Japan**

Shogo Ito, Kenta Yamauchi, and Hiroyuki Hioki

**Department of Cell Biology and Neuroscience, Juntendo University Graduate School of Medicine, Bunkyo-Ku, Tokyo 113-8421, Japan**

Shogo Ito, Kenta Yamauchi, Masato Koike, and Hiroyuki Hioki

**Laboratory for Cell Function Dynamics, RIKEN Center for Brain Science, Wako-City, Saitama, 351-0198, Japan**

Hiroshi Hama and Atsushi Miyawaki

**Advanced Research Institute for Health Sciences, Juntendo University, Bunkyo-Ku, Tokyo 113-8421, Japan**

Masato Koike

**Biotechnological Optics Research Team, RIKEN Center for Advanced Photonics, Wako-City, Saitama, 351-0198, Japan**

Atsushi Miyawaki

**Department of Multi-Scale Brain Structure Imaging, Juntendo University Graduate School of Medicine, Bunkyo-Ku, Tokyo 113-8421, Japan**

Hiroyuki Hioki

### Author contributions

S.I. contributed to conceptualization, methodology, investigation, formal analysis, writing – original draft, writing – review & editing and funding acquisition. K.Y. contributed to conceptualization, methodology, investigation, formal analysis, writing – original draft, writing – review & editing, project administration and funding acquisition. H.Hama contributed to investigation and writing – review & editing. M.K. contributed to writing – review & editing and funding acquisition. A.M. contributed to writing – review & editing and funding acquisition. H.Hioki contributed to conceptualization, investigation, writing – review & editing, project administration and funding acquisition.

### Corresponding authors

Kenta Yamauchi and Hiroyuki Hioki.

## Ethics declarations

### Competing interests

The authors declare that they have no competing interests.

## Notes

### Competing Interest Statement

The authors have declared no competing interest.

