## Supplementary Figures and Table. for "Plaque-associated endogenous IgG and its impact on immunohistochemical detection of mouse monoclonal IgG antibodies in mouse models of Alzheimer’s disease"

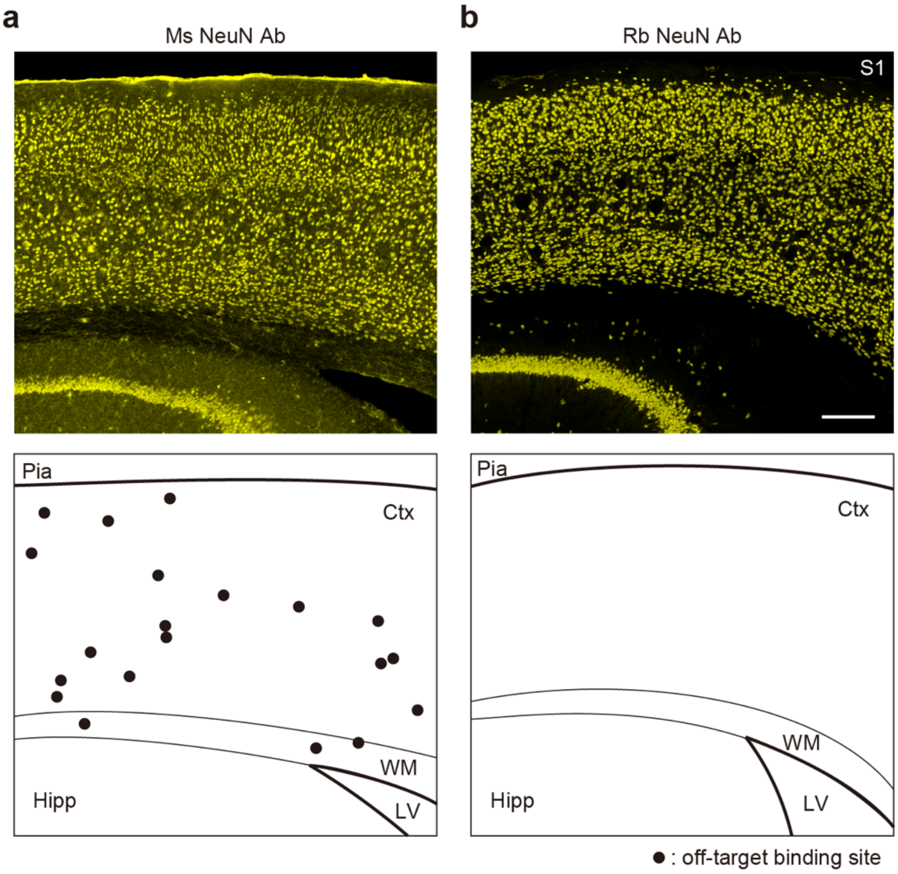


**Figure S1. Distribution of off-target binding in indirect IF using Ms and Rb NeuN IgG Abs in the S1 of *App*$^{NL\text{-}G\text{-}F/NL\text{-}G\text{-}F}$ mice.**

(**a**, **b**) Indirect IF using Ms (**a**) and Rb (**b**) NeuN Abs on brain sections of *App*$^{NL\text{-}G\text{-}F/NL\text{-}G\text{-}F}$ mice (yellow) (top panels). AF 488 conjugated Dk anti-Ms and Rb IgG (H+L) Abs are used for detection (n = 3 mice for each condition). The sites of off-target binding are plotted in the bottom panels (black circles). Hipp: hippocampus, LV: lateral ventricle, Pia: pia mater, S1: primary somatosensory cortex, and WM: white matter. Scale bar, 500 µm.

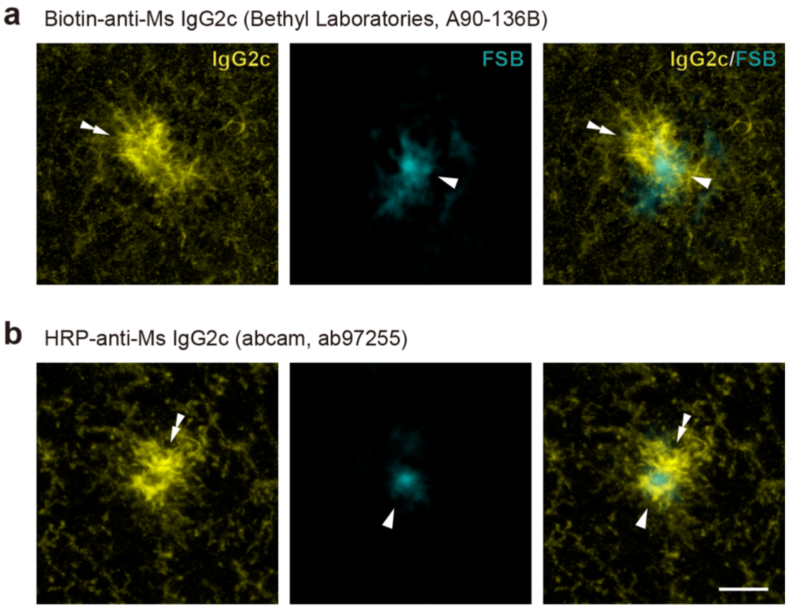


**Figure S2. Off-target binding of secondary Abs against Ms IgG2c on frozen brain sections of *App*$^{NL\text{-}G\text{-}F/NL\text{-}G\text{-}F}$ mice.**

(**a**, **b**) Applications of secondary Abs against Ms IgG2c conjugated with biotin (**a**) and HRP (**b**) (yellow) on brain sections of *App*$^{NL\text{-}G\text{-}F/NL\text{-}G\text{-}F}$ mice (n = 3 animals for each condition). Aβ plaques are labeled with FSB (cyan). Merged images of the secondary Abs and FSB are shown in the rightmost panels. Double arrowheads and arrowheads indicate off-target bindings of secondary Abs and Aβ plaques. Scale bar, 20 μm.

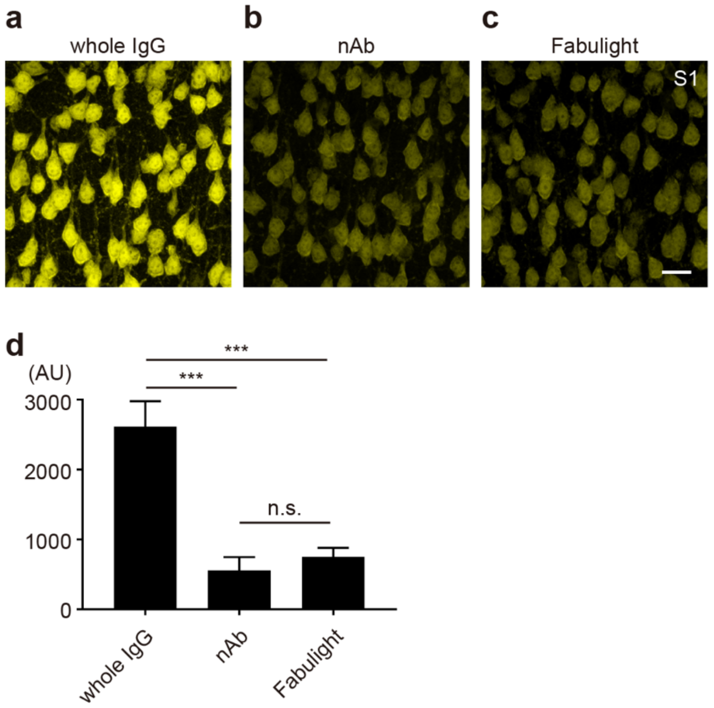


**Figure S3. NeuN IF using anti-Ms IgG1 secondary Abs with different Ab forms.** (**a-c**) NeuN IF in the S1 of wild-type mice using secondary Abs against Ms IgG1 with different Ab forms. A whole IgG Ab (**a**), nAb (**b**) and Fabulight Ab (Fab fragments of IgG) (**c**) conjugated with AF 488 are used for detection. (**d**) A histogram representing fluorescence signal intensities (AU) of NeuN positive cells in the layer2/3 of S1 (n = 492 cells, whole IgG Ab; n = 553 cells, nAb; n = 473 cells, Fabulight; n = 6 mice for each condition; $F = 124.8$, $df = 2$, $P < 0.0001$, ANOVA, ***$P < 0.0001$; Tukey's multiple comparisons test). Data are represented as means ± SDs. n.s.: not significant. Scale bar, 20 μm.

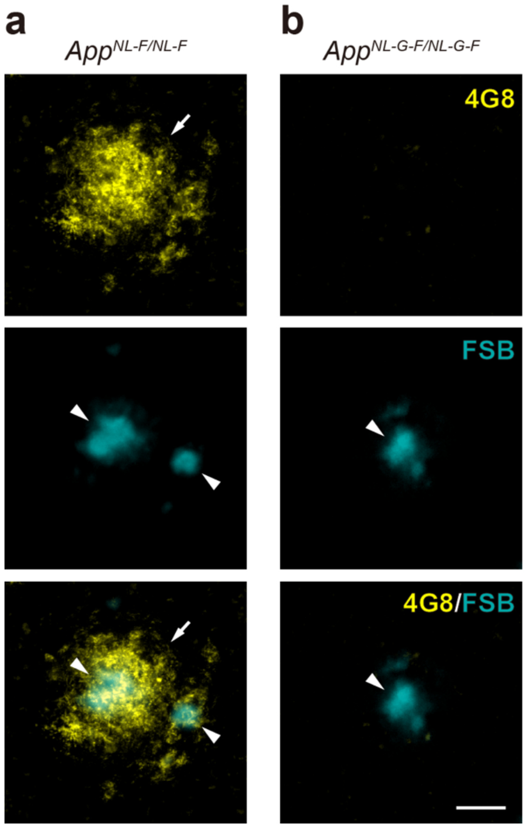


**Figure S4. Undetectable immunoreactivity of 4G8 Ab on frozen brain sections of** $\boldsymbol{App^{NL\text{-}G\text{-}F/NL\text{-}G\text{-}F}}$ **mice.**

(**a, b**) Direct IF using AF 488-conjugated 4G8 Ab (yellow) on brain sections of $App^{NL\text{-}F/NL\text{-}F}$ (**a**) and $App^{NL\text{-}G\text{-}F/NL\text{-}G\text{-}F}$ (**b**) mice (n = 3 mice for each condition). Aβ plaques are labeled with FSB (cyan). Merged images of the IF and FSB are shown in the bottommost panels. Arrows and arrowheads indicate signals of 4G8 Ab and FSB, respectively. Scale bar, 20 µm.

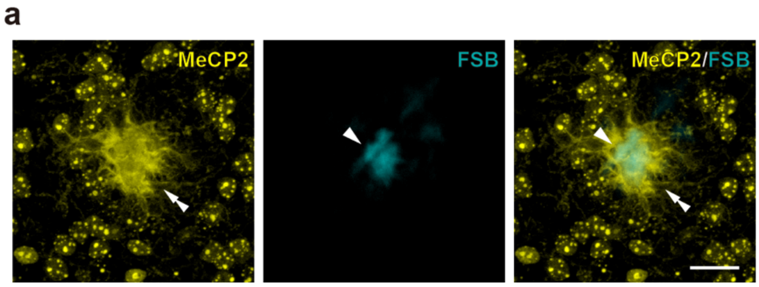


**Figure S5. Indirect IF for MeCP2 on frozen brain sections of *App*$^{NL\text{-}G\text{-}F/NL\text{-}G\text{-}F}$ mice with a secondary Ab against Ms IgG2c.**

(**a**) Indirect IF with Ms monoclonal IgG2c Ab against MeCP2 on frozen brain sections of *App*$^{NL\text{-}G\text{-}F/NL\text{-}G\text{-}F}$ mice (n = 3 mice for each condition). A secondary Ab against Ms IgG2c is used for the detection (yellow). Off-target binding (double arrowheads) to PA-IgG is visible among MeCP2 immunosignals (cell nuclei with bright foci). Aβ plaques are labeled with FSB (arrowheads, cyan). Merged image of the secondary Ab and FSB is shown in the rightmost panels. Scale bar, 20 μm.

Supplementary Table 1: Genotyping primers used in this study.

| *Allele* | *Primer name* | *Primer sequence (5’-3’)* |
|---|---|---|
| $App^{NL-F}$ | *E16WT* | *ATCTCGGAAGTGAAGATG* |
| $App^{NL-F}$ | *WT* | *TGTAGATGAGAACTTAAC* |
| $App^{NL-F}$ | *E16MT* | *ATCTCGGAAGTGAATCTA* |
| $App^{NL-F}$ | *LoxP* | *CGTATAATGTATGCTATACGAAG* |
| $App^{NL-G-F}$ | *common_0* | *CTCCTTGTGGCTGGCGGTCACAC* |
| $App^{NL-G-F}$ | *common_1* | *CTATCGTGGACCGAGAATGGTCATG* |
